# Long-term temporal trends in gastrointestinal parasite infection in wild Soay sheep

**DOI:** 10.1101/2022.04.28.489843

**Authors:** Adam D. Hayward, Jerzy M. Behnke, Dylan Z. Childs, Yolanda Corripio-Miyar, Andy Fenton, Mariecia D. Fraser, Fiona Kenyon, Tom N. McNeilly, Robin J. Pakeman, Amy B. Pedersen, Josephine M. Pemberton, Amy R. Sweeny, Ken Wilson, Jill G. Pilkington

## Abstract

Monitoring the prevalence and abundance of parasites over time is important for addressing their potential impact on host life-histories, immunological profiles, and their influence as a selective force. Only long-term ecological studies have the potential to shed light on both the temporal trends in infection prevalence and abundance and the drivers of such trends, because of their ability to dissect drivers that may be confounded over shorter time scales. Despite this, only a relatively small number of such studies exist. Here, we analysed changes in the prevalence and abundance of gastrointestinal parasites in the wild Soay sheep population of St Kilda across 31 years. The host population density has increased across the study, and population density is known to increase parasite transmission, but we found that density and year explained temporal variation in parasite prevalence and abundance independently. Prevalence of both strongyle nematodes and coccidian microparasites increased during the study, and this effect varied between lambs, yearlings and adults. Meanwhile, abundance of strongyles was more strongly linked to host density than to temporal (yearly) dynamics, while abundance of coccidia showed a strong temporal trend without any influence of density. Strikingly, coccidian abundance increased threefold across the course of the study in lambs, while increases in yearlings and adults were negligible. Our decades-long, intensive, individual-based study will enable the role of environmental change and selection pressures in driving these dynamics to be determined, potentially providing unparalleled insight into the drivers of temporal variation in parasite dynamics in the wild.

**Key findings:**

- We studied temporal trends in gastrointestinal parasites of wild sheep over 31 years
- Year and host population density explained temporal variation in parasites independently
- Prevalence of both strongyle nematodes and coccidia varied across the study period
- Abundance of strongyles was more closely linked to host density than year
- Abundance of coccidia increased threefold over time in lambs, but did not vary in adults

## Introduction

Parasite infections are widely known to follow seasonal patterns, with variation in prevalence and abundance of parasites apparent within each year (Altizer *et al*., 2006). This seasonality is known to be driven by a number of factors, including climatic effects on parasite development and mortality, host physiology or behaviour, or seasonal variability in the availability of naïve hosts (Cizauskas *et al*., 2015; Halvorsen *et al*., 1999; Haukisalmi & Henttonen, 1990; Haukisalmi *et al*., 1988; Montgomery & Montgomery, 1988; Pelletier *et al*., 2005; Rossin *et al*., 2010; Turner & Getz, 2010). Much less well understood is how parasite communities change across years, and specifically the extent to which parasites show long-term temporal trends in prevalence (the proportion of the host population that they infect) or abundance (the number of individual parasites harboured per host). Long-term studies of wild animal populations have frequently found evidence for variation and trends in population size across time (Andreassen *et al*., 2021; Armitage, 2012; Clutton-Brock *et al*., 2004;Crawley *et al*., 2021), and trends in phenological traits such as timing of breeding (Bonnet *et al*., 2019; Charmantier *et al*., 2008), morphological traits (Ozgul *et al*., 2009), and sexually-selected traits (Evans & Gustafsson, 2017). Given these long-term trends in host dynamics, one would expect similar temporal effects to apply to their parasites, but the link between long-term host and parasite dynamics is not well-understood. Similar long-term monitoring of the prevalence and abundance of parasites is important for addressing their potential impact on host life-histories, immunological profiles, and their influence as a selective force on host populations (Babayan *et al*., 2018; Hudson *et al*., 1998). Such long-term changes may be linked to a number of drivers, including: climatic variation (Haukisalmi & Henttonen, 1990; Huntley *et al*., 2014), since the development of transmission stages and vectors may be temperature- and moisture-dependent (Nordenfors *et al*., 1999; O’Connor *et al*., 2006; Rose *et al*., 2014); changes in host demography, for example if more vulnerable host age or sex classes change in relative abundance; or changes in selection on host immune function and/or related traits. Only long-term studies offer the power to identify temporal trends in parasite prevalence and abundance and determine their drivers, and indeed the power of studies to make inferences about the impact of diseases on host health, fitness and population dynamics is positively associated with their duration (Barroso *et al*., 2021).

The prevalence and abundance of many common and rare parasite species has been shown to be remarkably stable across years in studies of wild rodent populations (Bajer *et al*., 2004; Behnke *et al*., 2018; Behnke *et al*., 2008; Haukisalmi *et al*., 1988; Knowles *et al*., 2013). Where temporal variation does occur, it appears to consist of fluctuations between years rather than consistent trends across time (Behnke *et al*., 1999; Grzybek *et al*., 2015; Haukisalmi & Henttonen, 2000; Irvine *et al*., 2000; Lachish *et al*., 2011; Peet *et al*., 2022). In some cases, however, both colonisation of hosts by novel species and extinction of previously prevalent species have been reported (Grzybek *et al*., 2015; Kennedy *et al*., 2001; Lyndon & Kennedy, 2001). The majority of these studies take place across a maximum of 10 years, and use destructive sampling of a relatively small number of hosts in order to identify and enumerate the species present. Two recent studies have, however, found evidence for consistent temporal trends in parasite prevalence and abundance. In a 26-year study of the helminth fauna of a population of wood mice *(Apodemus sylvaticus)* in NE England, the parasite community was dominated by the nematode *Heligmosomoides polygyrus*, which showed a marked decline in prevalence and abundance across the study period, patterns that were not observed for other parasite species in the system (Behnke *et al*., 2021). Further, a similar decline in *H. polygyrus* faecal egg count was observed over 6 years in *A. sylvaticus* populations in NW England (Sweeny *et al*., 2021). These studies may have been able to detect these temporal trends because of their sampling regimes, which occurred over a long time scale (Behnke *et al*., 2021) or were non-destructive (Sweeny *et al*., 2021), both leading to a larger sample size and hence potentially greater power to detect consistent temporal trends. Perhaps most importantly, both studies returned to the same sampling sites every year, while many destructive sampling studies sample intermittently, leaving gaps in the data in order to enable host populations to recover. The lack of evidence for temporal trends may therefore stem from a combination of relatively small sample sizes, short time spans and intermittent sampling. Studies sampling sufficient individuals consistently at the same sites across a longer period would therefore seem to offer the greatest ability to discern temporal variation.

In addition to changes in the prevalence and mean abundance of parasites, temporal changes in the distribution of parasites among individuals across time may also be important (Grenfell *et al*., 1995; Haukisalmi & Henttonen, 1999). Generally, only hosts with the highest parasite burdens will experience fitness effects (although tolerance of infection may play a role in mitigating this), and the number and proportion of hosts to which this applies will depend on the distribution of parasites across the host population (Poulin & Vickery, 1993). Parasites tend to follow a negative binomial distribution among hosts, the shape of which dictates that the majority of hosts have relatively few parasites and a long ‘tail’ of a few hosts with high parasite burdens (Haukisalmi, 1986; Wilson *et al*., 2002; Woolhouse *et al*., 1997). The negative binomial distribution is characterised by the aggregation parameter *k*, and as *k* increases, the distribution becomes essentially Poisson or random. Thus, as *k* increases a greater proportion of the hosts fall into the tail of the distribution with high burden and parasites may be more likely to act as a regulatory factor for hosts (Anderson & May, 1978; May & Anderson, 1978). Variation in *k* across time may be generated through a number of processes that may serve to reduce aggregation, such as parasite mortality and parasite-induced host mortality, and increase it, such as heterogeneity in host resistance to infection as a result of genes or the environment (Anderson & Gordon, 1982). Determining if, for example, parasite distributions are becoming more or less aggregated across time may help us to determine which of these processes are occurring in a given host-parasite system, and lead us to identify the ecological or evolutionary drivers.

In this study, we determined how parasite prevalence and abundance have changed over 31 years in the wild Soay sheep population living in the St Kilda archipelago, NW Scotland.

Since the study of this population began in 1985, longitudinal data have been collected on the prevalence and abundance of several gastrointestinal parasite species, including apicomplexan and helminth parasites, with an abundance of associated data on host phenotype and environmental conditions (Clutton-Brock & Pemberton, 2004b). The sheep population has experienced significant temporal change during this time, with average body size declining (Ozgul *et al*., 2009) and population size increasing along with grassland productivity (Crawley *et al*., 2021). The known impacts of gastrointestinal parasites on condition and fitness in the Soay sheep (Craig *et al*., 2008; Gulland, 1992; Hayward *et al*., 2011) suggest that parasites may play a role in these dynamics, and the long-term nature of the study and abundance of metadata make the Soay sheep an excellent system in which to study long-term temporal trends in parasite infection and its drivers. Here, we take the first step by describing temporal changes in four parasite taxa across 31 years.

## Materials and methods

### Study population and data collection

We collected data from the wild population of Soay sheep living on Hirta in the St Kilda archipelago, NW Scotland (57°49’N 08°34’W). The population are descendants of early European domestic sheep that were introduced to the island of Soay several thousand years ago (Clutton-Brock & Pemberton, 2004a); in 1934-5, 108 sheep were moved from Soay onto Hirta and the population has since expanded across the island. The population living in the Village Bay area of Hirta has been intensively studied since 1985, with data collected on births, deaths, morphometrics, genetics, parasite infections and abundance, spatial use and environmental variation (Clutton-Brock & Pemberton, 2004a). Summer population density is estimated by ten censuses of the population in August of each year; any animal seen in any census is included in the population estimate (Fig. S1).

Since 1988, faecal samples have been collected and gastrointestinal parasite infection quantified as faecal egg counts (FEC), using a modified version of the McMaster egg-counting technique, accurate to 100 eggs per gram (epg) of faeces (M.A.F.F., 1986). Around 85% of the parasite data included in this study was collected by one of the authors (JGP). The McMaster egg-counting technique has been shown to be a good index of actual parasite burden in Soay sheep, both on St Kilda and elsewhere (Wilson *et al*., 2004). The present study exploits data collected in August of each year when animals were captured for weighing, skeletal measurement and blood and faecal sampling. Faecal samples were collected *per rectum* and stored at 4°C until processing, usually within a week of collection, all of which took place on Hirta. The parasite fauna in Soay sheep is very similar to that of domestic sheep on the mainland, with the notable exception of the nematode *Haemonchus contortus*. Among the principal taxa are strongyle nematodes, a group of six species of the gastrointestinal tract, namely *Teladorsagia circumcincta, Trichostrongylus axei*, *Trichostrongylus vitrinus*, *Chabertia ovina*, *Bunostomum trigonocephalum*, and *Strongyloides papillosus*. Since the eggs of these species are indistinguishable by eye, they are counted as a single ‘strongyle’ FEC. The sheep also experience infection with 11 species of apicomplexan microparasites of the genus *Eimeria*, the oocysts of which are again difficult to distinguish by eye and which are therefore incorporated as a single ‘coccidia’ faecal oocyst count (FOC). Strongyles and coccidia are by far the most prevalent and abundant parasite taxa present on St Kilda (Craig, 2005). In addition, eggs of the nematodes *Nematodirus* spp, *Trichuris ovis*, and *Capillaria longipes* are enumerated, and the presence or absence of eggs of the cestode *Moniezia expansa* is recorded. Data on strongyles were collected 1988-2018, while collection for the other taxa commenced in 1993; the exceptionally low prevalence (<1%) of *T. ovis* and *C. longipes* meant that we did not include them in any of our analyses (Table 1). Across the history of the project, several experiments have given anthelminthic treatments to a small number of animals in order to determine the effects on parasite burden and particularly survival and reproduction (Craig *et al*., 2009; Gulland, 1992). Most of these were long-lasting intraruminal boluses that released albendazole across several months and which were shown to be associated with lower worm burden at death in the following spring (Craig *et al*., 2009). Our analyses aimed to quantify changes in the annual prevalence of four of these parasite taxa (strongyles, coccidia, *Nematodirus* spp., *M. expansa)*, and abundance of strongyles and coccidia across the 31 years of the study.

**Table 1.** Number of samples (N), host individuals (IDs) and summary statistics for the parasite taxa analysed. The mean abundance of *M. expansa* is NA because only presence or absence is noted.

| Parasite | N | IDs | Prevalence | Mean abundance | SD abundance |
| --- | --- | --- | --- | --- | --- |
| Strongyles | 6873 | 3326 | 0.739 | 392.35 | 836.84 |
| Coccidia |  |  | 0.912 | 4365.52 | 12971.89 |
| *Nematodirus* spp. |  |  | 0.125 | 41.25 | 146.85 |
| *M. expansa* | 6077 | 3008 | 0.091 | NA | NA |
| *T. ovis* |  |  | 0.005 | 0.68 | 11.96 |
| *C. longipes* |  |  | 0.007 | 0.81 | 10.15 |

### Statistical analyses

All statistical analyses were conducted using R ver. 4.1.2 (R Core Team, 2021).

For each parasite group (strongyles, coccidia, *Nematodirus*, *M. expansa*), we first modelled changes in prevalence across time. We fitted generalized additive mixed-effects models (GAMMs) using the R package ‘gamm4’ (Wood & Scheipl, 2020). GAMMs fit non-parametric smoothing terms for continuous explanatory variables, meaning that the response variable is not restricted to following parametric forms (e.g. linear, quadratic). To determine the temporal dynamics of infection, we fitted binomial models, such that each sample scored 0 if the parasite was absent and 1 if it was present. We then tested a series of models to determine how prevalence changed across the years of the study, and whether changes in prevalence differed between age or sex classes. All models included individual animal identity and year as random effects in order to account for repeated sampling of individuals across years. We also included a categorical fixed effect (“Treatment”) in each model to account for anthelminthic treatments that were given as part of experiments in some years of the long-term study. Animals given an anthelminthic treatment in the year prior to data collection were scored as 1, while animals that did not received a 0; only 133/6873 samples (~2%) were associated with a treatment in our data set. We took this conservative approach because of the long-lasting effect of treatments on worm burden (Craig *et al*., 2009). All models also included a 3-level categorical fixed effect for age group, distinguishing lambs (animals aged ~4 months) from yearlings (animals aged ~16 months) and adults (animals aged ~28 months and older), due to the generally higher prevalence and abundance of parasites in younger animals (Wilson *et al*., 2004); 37% of our data was from lambs, 15% from yearlings and 48% from adults. All models included sex as a fixed categorical variable with two levels (67% female; 33% male), and the interaction between age class and sex.

To this ‘base’ model, we then added fixed effects describing temporal trends as described in detail in Table 2. Given the increase in population density (PD) across time (Fig. S1), and the role of host density in driving parasite transmission, accounting for PD in our analyses was essential. In models 1-8, we explored main effects of both year and PD, fitting models with linear and/or non-parametric smooth terms for year and/or PD. Linear models tested for directional trends, while smooth terms tested for non-linear variation across time. In all models, year and PD were both standardised to mean = 0 and SD = 1 in order to aid model convergence. In models 9-23, we then introduced interactions between smooth terms for year and/or PD with age class, sex, or the combined age/sex category; we used smooth rather than linear terms since the smooth simply reduces to a linear function if a linear association is present. Each of models 9-23 always included smooth terms for both year and PD, ensuring that any effects of either that we observed were independent of the other variable. We compared the AIC values for each model, and considered models with lower AIC values to be better supported by the data. Models within 2 AIC units of the best model were considered to receive some support from the data, and this is reflected in our presentation of results.

**Table 2.** A comparison of models describing temporal trends in the prevalence of strongyles and coccidia in St Kilda Soay sheep. In the model terms list, “s(Year/PD)” indicates a non-parametric smooth function was fitted to the variable inside parentheses. Note that both year and PD (population density) were standardized to mean = 0 and SD = 1 to aid model convergence. “ΔAIC” is the difference in AIC value between the model in question and the model with the lowest AIC value, which is highlighted in bold. Models shaded in grey fall within AIC = 2 of the best-supported model and are considered to receive some support from the data.

| Model | Terms | Strongyles |  | Coccidia |  |
| --- | --- | --- | --- | --- | --- |
| | | AIC | ΔAIC | AIC | ΔAIC |
| 0 | Null | 6126.42 | 35.53 | 2675.48 | 0.88 |
| 1 | Year | 6116.49 | 25.60 | **2674.60** | **0.00** |
| 2 | PD | 6120.53 | 29.64 | 2676.88 | 2.28 |
| 3 | Year + PD | 6114.70 | 23.82 | 2676.18 | 1.57 |
| 4 | s(Year) | 6110.22 | 19.33 | 2676.60 | 1.99 |
| 5 | s(PD) | 6116.23 | 25.34 | 2678.88 | 4.28 |
| 6 | s(Year) + PD | 6111.42 | 20.53 | 2678.18 | 3.58 |
| 7 | Year + s(PD) | 6112.02 | 21.13 | 2678.18 | 3.58 |
| 8 | s(Year) + s(PD) | 6107.92 | 17.03 | 2680.18 | 5.58 |
| 9 | s(Year):Age + s(PD) | 6100.26 | 9.37 | 2686.51 | 11.91 |
| 10 | s(Year):Sex + s(PD) | 6110.94 | 20.05 | 2684.08 | 9.48 |
| 11 | s(Year):Age:Sex + s(PD) | 6109.60 | 18.71 | 2695.03 | 20.42 |
| 12 | s(Year) + s(PD):Age | 6093.03 | 2.14 | 2687.33 | 12.73 |
| 13 | s(Year) + s(PD):Sex | 6100.42 | 9.53 | 2684.19 | 9.58 |
| 14 | s(Year) + s(PD):Age:Sex | 6096.67 | 5.78 | 2698.63 | 24.03 |
| 15 | s(Year):Age + s(PD):Age | **6090.89** | **0.00** | 2693.94 | 19.33 |
| 16 | s(Year):Age + s(PD):Sex | 6091.70 | 0.81 | 2690.53 | 15.93 |
| 17 | s(Year):Age + s(PD):Age:Sex | 6092.86 | 1.97 | 2705.15 | 30.55 |
| 18 | s(Year):Sex + s(PD):Age | 6096.98 | 6.09 | 2691.27 | 16.66 |
| 19 | s(Year):Sex + s(PD):Sex | 6105.20 | 14.31 | 2688.01 | 13.40 |
| 20 | s(Year):Sex + s(PD):Age:Sex | 6100.56 | 9.67 | 2702.47 | 27.87 |
| 21 | s(Year):Age:Sex + s(PD):Age | 6097.58 | 6.70 | 2702.54 | 27.94 |
| 22 | s(Year):Age:Sex + s(PD):Sex | 6103.05 | 12.16 | 2698.97 | 24.36 |
| 23 | s(Year):Age:Sex + s(PD):Age:Sex | 6098.77 | 7.88 | 2713.21 | 38.60 |

For strongyles and coccidia, we then repeated a similar analysis in order to identify temporal patterns of abundance measured as FEC and FOC respectively, once again accounting for PD. For both parasite groups, we used negative binomial models with a log link function in ‘gamm4’, where the variance is modelled as *σ*^2^ = *μ* + *μ*^2^/*k*, where *μ* is the mean and *k* is the dispersion parameter. The dispersion parameter was estimated from the data and provided to the model as a starting value. We fitted the same 23 models as described above for prevalence and as shown in Table 3.

**Table 3.** A comparison of models describing temporal trends in the abundance of strongyles and coccidia in St Kilda Soay sheep. In the model terms list, “s(Year/PD)” indicates a non-parametric smooth function was fitted to the variable inside parentheses. Note that both year and PD (population density) were standardized to mean = 0 and SD = 1 to aid model convergence. “ΔAIC” is the difference in AIC value between the model in question and the model with the lowest AIC value, which is highlighted in bold.

| Model | Terms | Strongyles |  | Coccidia |  |
| --- | --- | --- | --- | --- | --- |
| | | AIC | ΔAIC | AIC | ΔAIC |
| 0 | Null | 85374.56 | 16.50 | 104487.40 | 28.90 |
| 1 | Year | 85370.87 | 12.81 | 104476.60 | 18.10 |
| 2 | PD | 85363.82 | 5.76 | 104489.40 | 30.90 |
| 3 | Year + PD | 85363.89 | 5.83 | 104478.20 | 19.70 |
| 4 | s(Year) | 85370.98 | 12.92 | 104478.60 | 20.10 |
| 5 | s(PD) | 85364.02 | 5.96 | 104491.40 | 32.90 |
| 6 | s(Year) + PD | 85365.89 | 7.83 | 104480.20 | 21.70 |
| 7 | Year + s(PD) | 85365.00 | 6.94 | 104480.20 | 21.70 |
| 8 | s(Year) + s(PD) | 85367.00 | 8.94 | 104482.20 | 23.70 |
| 9 | s(Year):Age + s(PD) | 85371.17 | 13.11 | 104464.80 | 6.30 |
| 10 | s(Year):Sex + s(PD) | 85369.77 | 11.71 | 104482.90 | 24.40 |
| 11 | s(Year):Age:Sex + s(PD) | 85382.67 | 24.61 | **104458.50** | **0.00** |
| 12 | s(Year) + s(PD):Age | 85360.61 | 2.55 | 104485.00 | 26.50 |
| 13 | s(Year) + s(PD):Sex | 85360.95 | 2.89 | 104483.00 | 24.50 |
| 14 | s(Year) + s(PD):Age:Sex | **85358.06** | **0.00** | 104496.40 | 37.90 |
| 15 | s(Year):Age + s(PD):Age | 85369.26 | 11.20 | 104468.90 | 10.40 |
| 16 | s(Year):Age + s(PD):Sex | 85368.43 | 10.37 | 104469.90 | 11.40 |
| 17 | s(Year):Age + s(PD):Age:Sex | 85366.66 | 8.60 | 104480.40 | 21.90 |
| 18 | s(Year):Sex + s(PD):Age | 85366.18 | 8.12 | 104489.60 | 31.10 |
| 19 | s(Year):Sex + s(PD):Sex | 85366.95 | 8.89 | 104487.00 | 28.50 |
| 20 | s(Year):Sex + s(PD):Age:Sex | 85364.06 | 6.00 | 104500.50 | 42.00 |
| 21 | s(Year):Age:Sex + s(PD):Age | 85380.51 | 22.45 | 104462.70 | 4.20 |
| 22 | s(Year):Age:Sex + s(PD):Sex | 85380.32 | 22.26 | 104463.40 | 4.90 |
| 23 | s(Year):Age:Sex + s(PD):Age:Sex | 85377.88 | 19.82 | 104474.40 | 15.90 |

Finally, we tested for changes in three parameters describing the distribution of strongyle FEC and coccidian FOC across time. For each year, we calculated the variance (*σ*^2^ = ∑_*i*_ (*x_i_* – *μ*)^2^/(*n* – 1)), the dispersion index (*I* = *σ*^2^/*μ*) and the negative binomial dispersion parameter (*k* = *μ*^2^/*σ*^2^ – *μ*), where *x_i_*, *μ* and *σ* are the value for an individual sample *i*, the population mean and the population standard deviation of FEC/FOC respectively, for that year. We then fitted a linear model testing for linear and quadratic changes in each of these parameters across time, accounting for PD. We then calculated age class-specific (lamb, yearling, adult) parameters and tested whether these differed between age classes in their distribution parameters across time, and then did the same for sex-specific parameters. All associations were tested with Wald F-tests.

## Results

### Temporal variation in prevalence

The prevalence of strongyles was influenced by year and population density (PD) independently. Of the main effect only models 0 to 8, the model with smooth terms for both year and PD had the lowest AIC (model 8 ΔAIC = −2.30 compared to model 4 with same terms but no smoothing, Table 2). The model with the lowest AIC overall was model 15, which fitted interactions between the smooth terms for both year and PD with age class (Table 2). The model predicted an approximately linear increase in the prevalence of strongyles in lambs and adults over time, with a curvilinear association in yearlings, predicting an increase in 1988-1995, a decline towards 2010 and then stasis thereafter (Figure 1A). In addition, the model predicted higher strongyle prevalence at higher host densities, although the shape of this association varied among age classes: the model predicted an approximately linear increase with PD in lambs, while in yearlings and adults the model predicted steep increases in prevalence from low to moderate PD and relative stasis at higher PD (Figure 1B). Models 16 and 17 had ΔAIC<2 and so were considered to receive some support; they both supported the interaction between year and age, but also provided support for interactions between PD and sex (model 16) and between PD, sex and age (model 17). These results are presented in Fig. S2.

**Figure 1.**
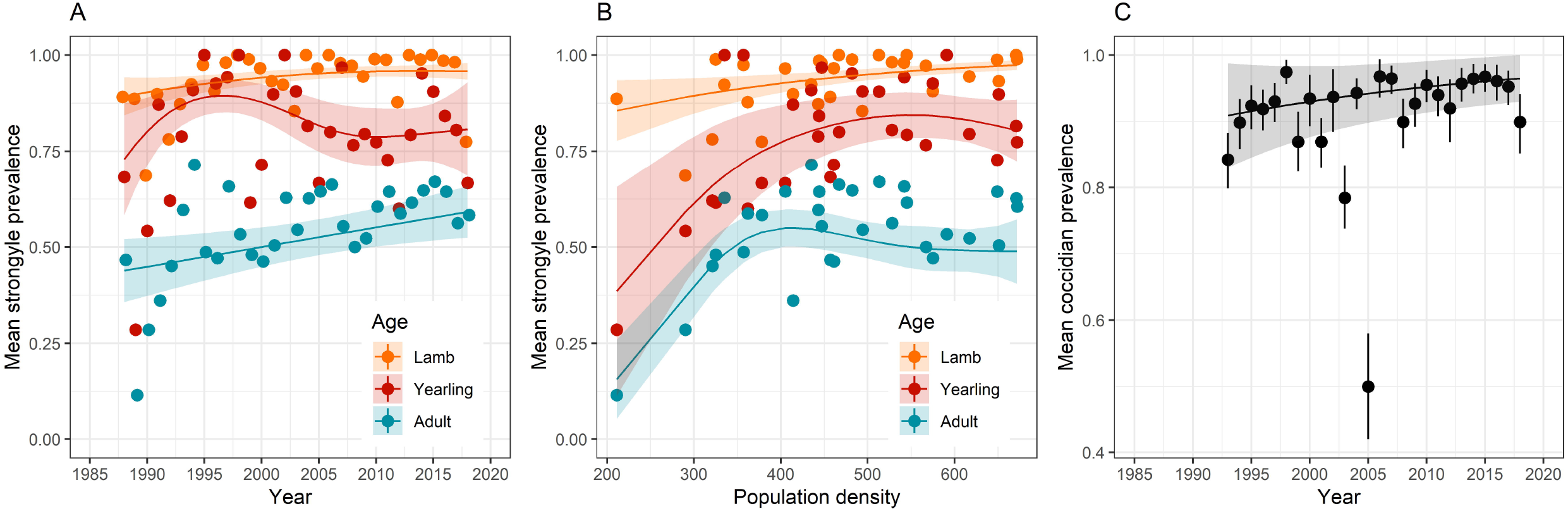
Variation in strongyle prevalence across both (A) the study period and (B) population density varied between age classes; (C) the prevalence of coccidia increased across the study period in an approximately linear fashion. Points show raw mean prevalence, while lines and shaded areas show predictions and standard errors from model 15 for strongyles and model 3 for coccidia in Table 2.

The prevalence of coccidia was only weakly influenced by year and PD. The model with the lowest AIC value was model 1, which fitted just a linear effect of year (Table 2). The null model, however, had a ΔAIC value of only +0.88, suggesting that the effect of year was only marginal. Also within ΔAIC of 2 were models 3 and 4 (Table 2), both of which included main effects of year. Models that included PD and interactions with age and sex fitted poorly. Overall, there appeared to be a weak increase in the prevalence of coccidia across the study period (Fig. 1C).

For both *Nematodirus* spp. and *M. expansa*, the best-supported model was the null model, suggesting no strong influence of either year or PD (Table S1). Although the prevalence of both varied between around 0.05 and 0.20 across the study period, neither linear trends nor non-parametric smooth terms were supported (Fig. S3). As described above, both *T. ovis* and *C. longipes* had mean prevalence of <1% and were not analysed (Figure S3).

### Temporal variation in abundance

Strongyle FEC was largely associated with population density rather than year (Table 3). For example, while models including year (e.g. models 1 and 4) generally had lower AIC than the null model, this was not true once PD was added to the model (e.g. compare models 5 and 8). The best-supported model, with AIC −2.55 compared to the next-best model, suggested that strongyle FEC varied with PD in a manner that varied between age and sex classes (model 14). Broadly, males and younger animals had higher FEC than females and adults, and the increase in FEC with increasing PD was greatest in lambs, while in yearlings and adults there was a relatively weak positive association (Fig. 2A). There were also subtle differences among the sexes, which may explain why this model fitted better than the solely age-specific PD model 12 (Table 3). For example, among lambs, the difference between the sexes was greatest at higher OD, while along yearlings, the difference between the sexes was greater at lower PD (Figure 2A).

**Figure 2.**
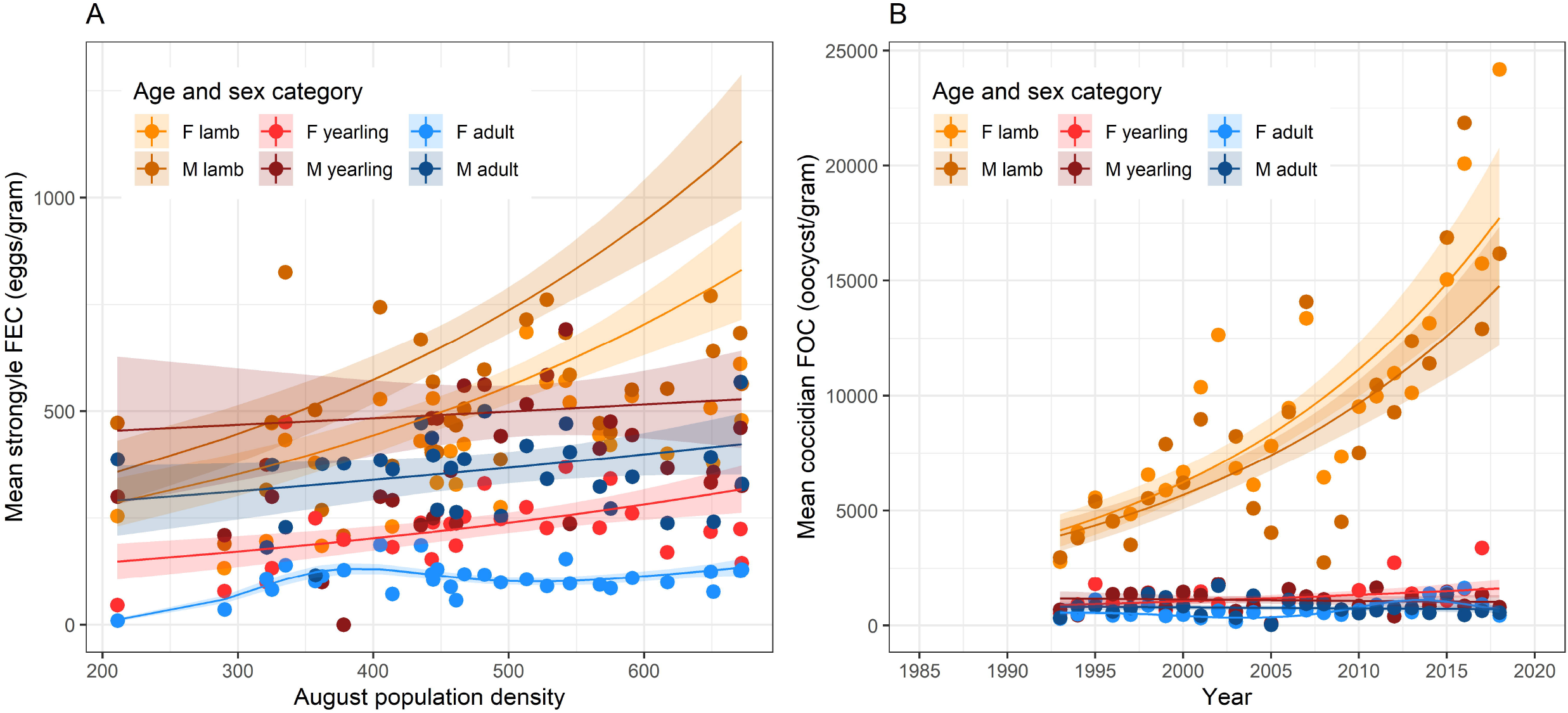
Generalized additive mixed-effects model-predicted variation in strongyle FEC and coccidian FOC from the best-supported models in Table 3. (A) Variation in strongyle FEC with population density in each of 6 age and sex classes; (B) variation in coccidian FOC across time in the same age and sex classes. Points show raw mean abundance, while lines and shaded areas show predictions and standard errors from model 14 for strongyles and model 11 for coccidia in Table 3.

In contrast to strongyle FEC, coccidian FOC was chiefly associated with year rather than population density (e.g. compare models 4, 5, and 8 in Table 3). The best-supported model (model 11) had AIC −4.20 relative to the next-best model and supported variation in coccidian FOC across years that varied between age and sex classes (Table 3). The most notable pattern was the increase in FOC among lambs, with the model predicting around a 3-fold increase in FOC across the course of the study (Fig. 2B). Meanwhile, FOC in yearlings and adults appears relatively stable across time. Looking more closely at the model predictions, however, sex differences become apparent, which explain why model 11 was best-supported. For example, the increase in FOC across time appears to have been slightly stronger in female lambs than in male lambs, while in yearlings and adults there appear to have been slight declines across time in males and slight increases in females, with a non-linear pattern apparent in adult females (Fig. S4).

### Abundance distribution parameters

None of the distributional parameters for strongyle FEC changed across time, and nor did changes across time vary between the sexes or age classes (Table S2; Figure 3A-C). However, all three distributional parameters for coccidian FOC varied across time. Variance in FOC increased with time (Table S3), but this increase was really only apparent in lambs (Figure 3D), matching the increase in mean abundance that we saw. The variance-to-mean ratio, also known as the dispersion parameter *I*, increased with time (Figure 3E), but this increase did not differ with age class or sex (Table S3). Finally, although the aggregation parameter *k* did not change with year as a main effect (Table S3), there was evidence for an interaction with age class, such that the aggregation parameter decreased with time in adults and especially yearlings but increased with time in lambs (Figure 3F).

**Figure 3.**
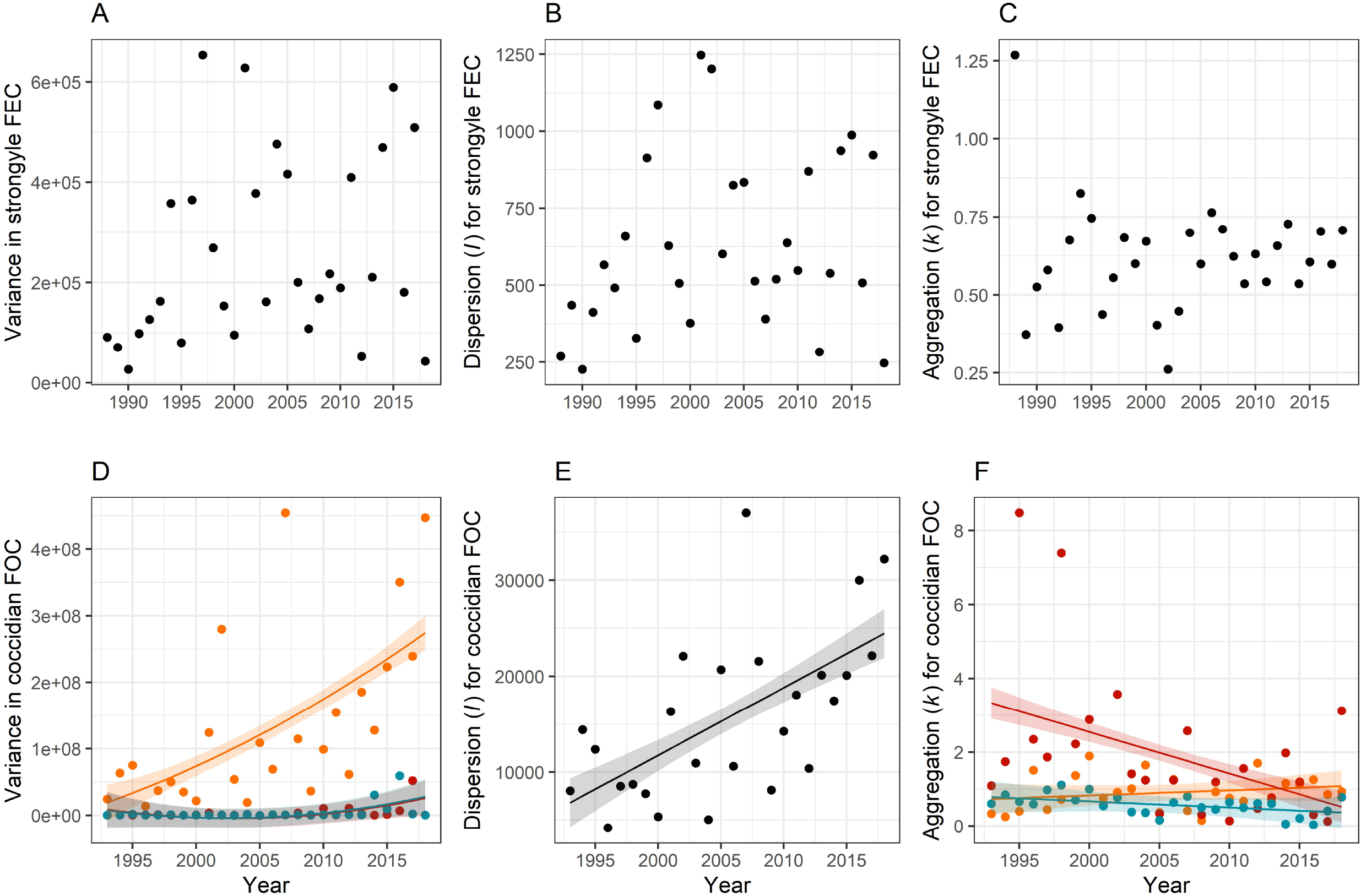
Temporal variation distribution parameters for (A – C) strongyle faecal egg count (FEC) and (D – F) coccidian faecal oocyst count (FOC). Points show raw parameters estimates from the data and lines show results of linear models ±1SE; lambs are in orange, yearlings in red, and adults in blue.

## Discussion

Long-term studies of wildlife disease are vital to enable us to understand how these systems respond to individual, population and environmental change, and to separate the drivers of host-pathogen/parasite dynamics (Barroso *et al*., 2021). In this study, we describe the nature of temporal trends in parasite infections in the Soay sheep of St Kilda across 31 years. Our results show that the prevalence and/or abundance of two prevalent and pathogenic types of parasite have changed across the study period in age- and sex-dependent ways.

Strongyles and coccidia are the most prevalent and abundant parasites infecting St Kilda Soay sheep, and here we show that the prevalence of both, and the abundance of coccidia, increased across time. These increases were over-and-above any effect of host population density, which was positively associated with prevalence and abundance of most of our parasite taxa, and which itself increased across time by around 5 sheep per year between 1988 and 2018 (Fig. S1; linear regression of density on year estimate 4.95±2.25SE, F=4.85, P=0.036). Furthermore, we accounted for host age and sex in our models, so these year-on-year trends are independent of these factors. Previous studies of wildlife disease have determined that changes in host population density can play a role in generating temporal patterns of disease prevalence, such as the case of tuberculosis in European badgers (*Meles meles*), red deer (*Cervus elaphus*), fallow deer (*Dama dama*) and wild boar (*Sus scrofa*), which has increased across time in these populations at least partly in response to host density (Barroso *et al*., 2020; Delahay *et al*., 2013; Vicente *et al*., 2013). Due to the particularly long-term nature of our study, we were able to disentangle effects of density and year in our analyses, and found that both influenced prevalence and/or abundance of strongyles and/or coccidia in an age-dependent manner. While prevalence of coccidia increased with time in a roughly linear fashion, prevalence of strongyles increased linearly in lambs and adults, but followed a non-linear trajectory in yearlings, with an increase in prevalence early in the study followed by stasis thereafter. Meanwhile, although prevalence of strongyles was high in lambs (94% overall), it still increased with host population density in a linear fashion; in yearlings and adults, the pattern was more of an increase between low and moderate density and stasis at higher density. Transmission is expected to be higher at higher density in this population (Hayward *et al*., 2009) and lambs at 4 months are likely to be more susceptible to infection due to their less effective parasite-specific immune responses compared to adults and yearlings (Sparks *et al*., 2019). The linear association between density and prevalence in lambs may reflect the fact that naïve lambs act as “sentinels” of infection pressure (Hayward *et al*., 2014), but once transmission reaches a certain level in older animals, their acquired immunity may prevent further increases in infection. This result was also reflected in that for strongyle abundance: lambs experienced increased strongyle FEC at higher density, while the effect of density was much weaker in adults and yearlings (Figure 2A). By far the most striking result was the marked increase in coccidian FOC across the study period in lambs that was largely absent from yearlings and adults (Figure 2B). Coccidian parasites can cause considerable clinical signs and pathology in lambs from domestic sheep flocks. In these livestock systems, infection levels generally peak just after weaning (Chartier & Paraud, 2012), which is just before we sampled the St. Kilda Soay Sheep in August. If lambs also act as sentinels of coccidian infection, our results suggest that infection pressure applied by coccidia has increased across the course of the study independently of host density. Explaining the temporal trends we observed, and particularly the striking increase in coccidia abundance in lambs, remains a task for further, detailed, analysis, but several potential mechanisms could be involved.

When explaining long-term temporal trends in ecology, climate and specifically global warming naturally spring to mind (Huntley *et al*., 2014). Since the start of the Soay sheep study in 1985, mean temperature across all parts of the year on St Kilda has increased by ~0.03°C per year (Crawley *et al*., 2021). Both strongyles and coccidia are directly transmitted, and both temperature and moisture are associated with the survival and development of strongyle larvae (O’Connor *et al*., 2006; Rose *et al*., 2014) and coccidian oocysts (Makau *et al*., 2017; Waldenstedt *et al*., 2001) in the external environment. These observations are reflected in the associations between (for example) warmer temperatures and greater parasite abundances in other wild (Holand *et al*., 2019; Mennerat *et al*., 2019) and domestic animal populations (Kenyon *et al*., 2009; van Dijk *et al*., 2009). Indeed, a warming climate has been predicted to cause an increase in infection pressure by gastrointestinal nematodes in domestic sheep (Morgan & van Dijk, 2012; Rose *et al*., 2016; Rose *et al*., 2015; van Dijk *et al*., 2008). Projected changes in precipitation are also likely to play a role (O’Connor *et al*., 2006). The availability of long-term climate data on St Kilda (Crawley *et al*., 2021), means that the role of climate in driving temporal trends in parasitology can be thoroughly investigated in future analyses.

Another factor potentially explaining the temporal trends in parasite prevalence and abundance could be changes in selection pressure in the population. Higher strongyle burdens (in particular) are associated with reduced winter survival in Soay sheep, an association that is restricted to lambs and absent in adults (Gulland, 1992; Hayward *et al*., 2011). If selection has weakened such that lambs with less effective immune responses and higher parasite burdens are now more likely to survive, increases in parasite abundance may become apparent. This hypothesis is, however, somewhat troubled by the observation that, despite the much lower strongyle-specific antibody levels in lambs compared to adults (Sparks *et al*., 2019), strongyle-specific antibodies are associated with survival in adult females, but not in lambs (Sparks *et al*., 2018). Selection on August FEC in lambs does vary among years, with FEC being associated with survival in low-mortality years but not high-mortality years (Hayward *et al*., 2011). Temporal changes in patterns of selection have been observed in other studies of wild populations: warming temperature can explain a shift from selection for larger to selection for smaller forehead patch size in male collared flycatchers (*Ficedula albicollis)* (Evans & Gustafsson, 2017) and weakening of selection on hatch date in pied flycatchers *(Ficedula hypoleuca)* (Visser *et al*., 2015). Despite all of these findings, no formal analysis of temporal or environment-dependent variation in selection on immunoparasitological traits has been conducted on the Soay sheep population.

We saw no evidence for changes in the distribution of strongyles across time, which was perhaps unsurprising given that we found no change in mean abundance. We did, however, observe variation in all three distributional parameters for coccidia (Fig. 3). An intriguing result from the point of view of parasite epidemiology is the age-specific trend in the aggregation parameter *k* for coccidian FOC (**Fig. 3F**), which suggests that FOC is becoming less aggregated among lambs (increasing *k*) and more aggregated in adults and especially yearlings (decreasing *k*), although the trends in lambs and adults are relatively weak. This result broadly supports theory that predicts that aggregation should decrease as encounters with parasites (i.e. the force of infection) increases (Gourbière *et al*., 2015). Given that *k* increases (and aggregation decreases) with the mean, it is perhaps unsurprising that we saw this result in lambs, and visual inspection of the distribution of FOC in lambs suggests a year-on-year reduction in aggregation (Figure S5). Where parasites are abundant, increasing *k* places a greater proportion of hosts under potentially health-limiting parasite burdens (Poulin & Vickery, 1993), suggesting that lambs are increasingly facing selection pressures from coccidian infection. A major advantage of the Soay sheep system is that hypotheses about selection on parasite resistance can be tested, thanks to the detailed data on survival, reproduction and parasitology that have been collected across the decades (Clutton-Brock & Pemberton, 2004a), plus a cache of stored blood samples that has been interrogated for parasite-specific antibody responses (Froy *et al*., 2019; Hayward *et al*., 2014; Sparks *et al*., 2019).

The strength of our study lies in its long-term nature: data have been collected at the same time in every year since 1988, making it a rarity among studies of wildlife disease (Barroso *et al*., 2021). The longitudinal nature of the study has enabled us to identify temporal trends and to separate them from effects of host density and demography. The next challenge is to make use of the associated meta-data in order to identify the forces that are driving the changes we have observed. On the other hand, the main weakness of our study lies in the lack of parasite species-specificity in our data, which arises because of the difficulty of distinguishing individual parasite species based on egg/oocyst morphology alone. The focus has been on faecal sampling throughout the St. Kilda Soay sheep study largely because of its non-invasive nature. Previous work on the Soay sheep has identified helminth species based on adult worm morphology by necropsy of animals that died over winter (Craig *et al*., 2006) and *Eimeria* species through labour-intensive sporulation of oocysts and identification based on morphology (Craig *et al*., 2007). These studies have provided intriguing results: for example, while *Trichostrongylus axei* and *T. vitrinus* dominate the strongyle species in lambs, *Teladorsagia circumcincta* is much more common in adults (Craig *et al*., 2006); meanwhile, *Eimeria* species diversity is high in lambs but declines with age (Craig *et al*., 2007). It is currently impossible to determine which species are responsible for the temporal trends observed here, but elucidating this is important, because of the different locales of the parasites within the host, and their potentially different impacts on host health. Application of new meta-barcoding techniques to both nematodes (Avramenko *et al*., 2015) and coccidia (Heitlinger *et al*., 2017) could help in this regard.

Overall, our results provide evidence for temporal trends in the prevalence and abundance of fitness-limiting parasites across three decades in a wild ungulate population. These trends were generally stronger in the youngest and most susceptible animals in the population than in older animals, and were absent for rarer parasite species. Decades of intensive, individual-based study of this population, plus new data on parasite species identity, will enable the role of environmental change and selection pressures in driving these dynamics to be determined, potentially providing unrivalled insight into the drivers of temporal variation in parasite dynamics in the wild.

## Acknowledgements

Thanks to Dan Nussey for insightful comments on earlier drafts of the manuscript. We thank the National Trust for Scotland for permission to work on St. Kilda and QinetiQ, Eurest and Kilda Cruises for logistics and support. We thank I. Stevenson and many volunteers who have collected field data and samples and all those who have contributed to keeping the project going.

## Financial support

Field data collection has been supported principally by NERC over many years, with some funding from the Wellcome Trust. ADH is supported by a Moredun Foundation Research Fellowship.

## Notes

### Competing Interest Statement

The authors have declared no competing interest.

