## Supporting Information for "Long-term temporal trends in gastrointestinal parasite infection in wild Soay sheep"

**Items appear in the order which they appear in the main text.**

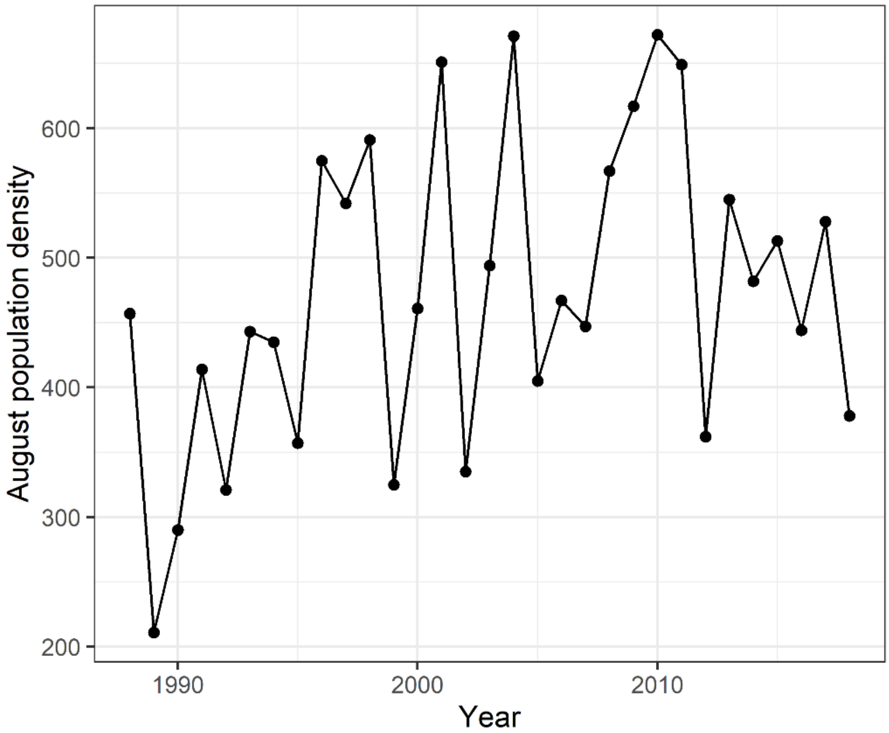


**Figure S1.** Village Bay population density, 1988-2018. A linear regression of population density on year suggests a significant increase of 4.95 (±2.25SE) sheep per year (F=4.85, P=0.036).

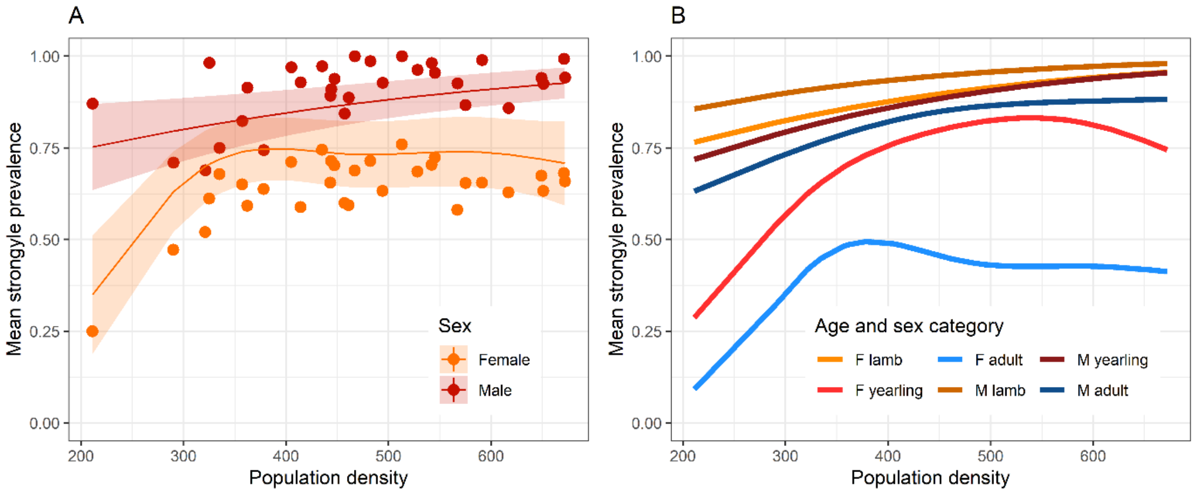


**Figure S2.** Variation in strongyle prevalence with population density varied between (A) the sexes according to model 16 (Table 2); and (B) a combination of age and sex category according to model 17 (Table 2). Points show raw mean prevalence, while lines and shaded areas show predictions and standard errors from models in Table 2. Neither model was as well-supported as model 15.

**Table S1.** A comparison of models describing temporal trends in the prevalence of *Nematodirus* spp and *Moniezia expansa* in St Kilda Soay sheep. In the model terms list, “s(Year/PD)” indicates a non-parametric smooth function was fitted to the variable inside parentheses. Note that both year and PD (population density) were standardized to mean = 0 and SD = 1 to aid model convergence. “ΔAIC” is the difference in AIC value between the model in question and the model with the lowest AIC value, which is highlighted in **bold**. Models shaded in grey fall within AIC = 2 of the best-supported model and are considered to receive some support from the data.

| | | ***Nematodirus*** | | ***Monezia*** | |
|---|---|---|---|---|---|
| **Model** | **Terms** | **AIC** | **ΔAIC** | **AIC** | **ΔAIC** |
| **0** | **Null** | **2874.19** | **0.00** | **3340.27** | **0.00** |
| 1 | Year | 2876.10 | 1.91 | 3341.84 | 1.57 |
| 2 | PD | 2874.27 | 0.09 | 3342.47 | 2.20 |
| 3 | Year + PD | 2876.08 | 1.90 | 3344.17 | 3.90 |
| 4 | s(Year) | 2878.10 | 3.91 | 3343.93 | 3.66 |
| 5 | s(PD) | 2876.27 | 2.09 | 3346.11 | 5.84 |
| 6 | s(Year) + PD | 2878.08 | 3.90 | 3348.82 | 8.55 |
| 7 | Year + s(PD) | 2878.08 | 3.90 | 3366.70 | 26.43 |
| 8 | s(Year) + s(PD) | 2880.08 | 5.90 | 3567.01 | NA |
| 9 | s(Year):Age + s(PD) | 2884.68 | 10.49 | 3357.76 | 17.49 |
| 10 | s(Year):Sex + s(PD) | 2881.37 | 7.18 | 3359.58 | 19.30 |
| 11 | s(Year):Age:Sex + s(PD) | 2897.02 | 22.84 | NA | NA |
| 12 | s(Year) + s(PD):Age | 2882.21 | 8.02 | 3571.84 | NA |
| 13 | s(Year) + s(PD):Sex | 2882.33 | 8.14 | 3370.66 | 30.39 |
| 14 | s(Year) + s(PD):Age:Sex | 2895.92 | 21.74 | NA | NA |
| 15 | s(Year):Age + s(PD):Age | 2885.14 | 10.95 | 3380.16 | NA |
| 16 | s(Year):Age + s(PD):Sex | 2886.98 | 12.80 | 3582.29 | NA |
| 17 | s(Year):Age + s(PD):Age:Sex | 2900.13 | 25.94 | NA | NA |
| 18 | s(Year):Sex + s(PD):Age | 2888.21 | 14.02 | 3579.50 | NA |
| 19 | s(Year):Sex + s(PD):Sex | 2883.25 | 9.06 | 3577.75 | NA |
| 20 | s(Year):Sex + s(PD):Age:Sex | 2896.86 | 22.67 | NA | NA |
| 21 | s(Year):Age:Sex + s(PD):Age | 2893.98 | 19.80 | NA | NA |
| 22 | s(Year):Age:Sex + s(PD):Sex | 2898.50 | 24.31 | NA | NA |
| 23 | s(Year):Age:Sex + s(PD):Age:Sex | 2912.18 | 37.99 | NA | NA |

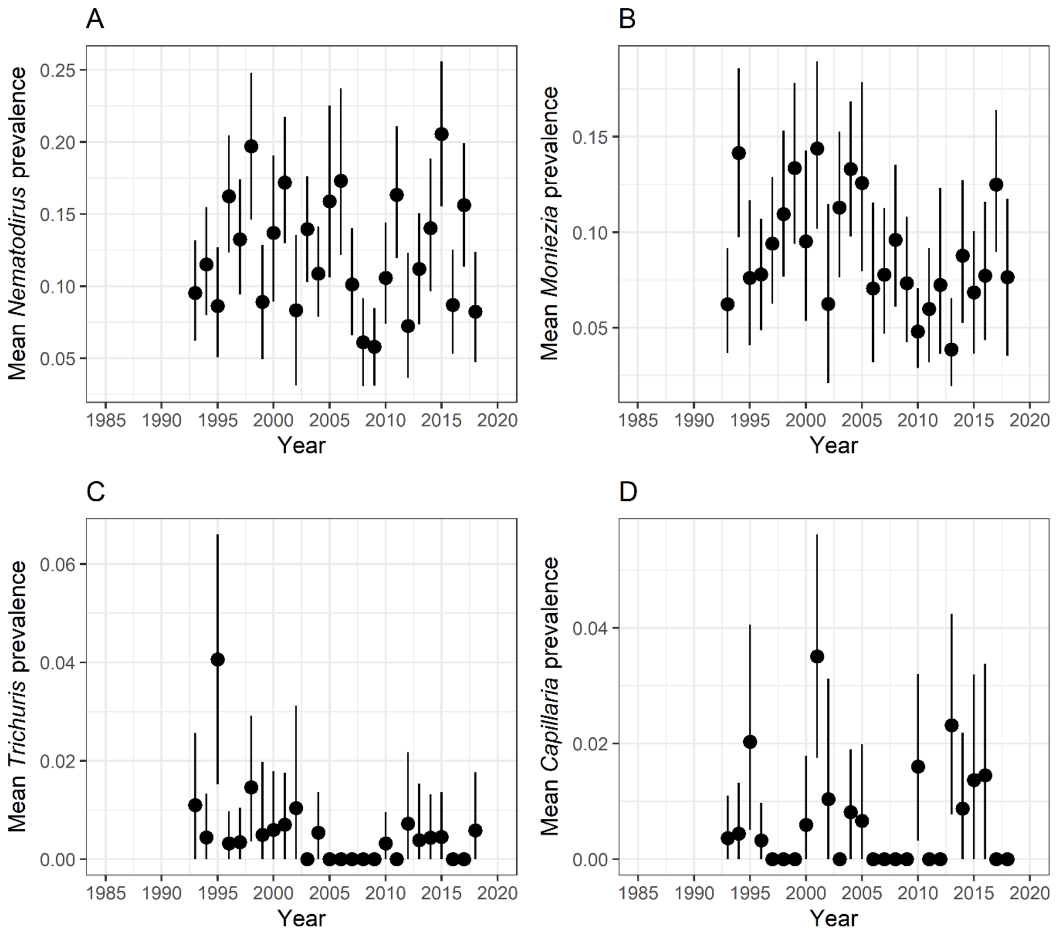


**Figure S3.** Variation in the prevalence of (A) *Nematodirus*; (B) *M. expansa*; (C) *T. ovis*; (D) *C. longipes* across the study period. Points show raw mean prevalence and standard errors. Neither linear nor non-parametric smooth terms were supported in analysis of *Nematodirus* and *M. expansa*, suggesting the absence of temporal trends for these parasites (Table S1). The prevalence of both *T. ovis* and *C. longipes* were <1% and so not considered for analysis.

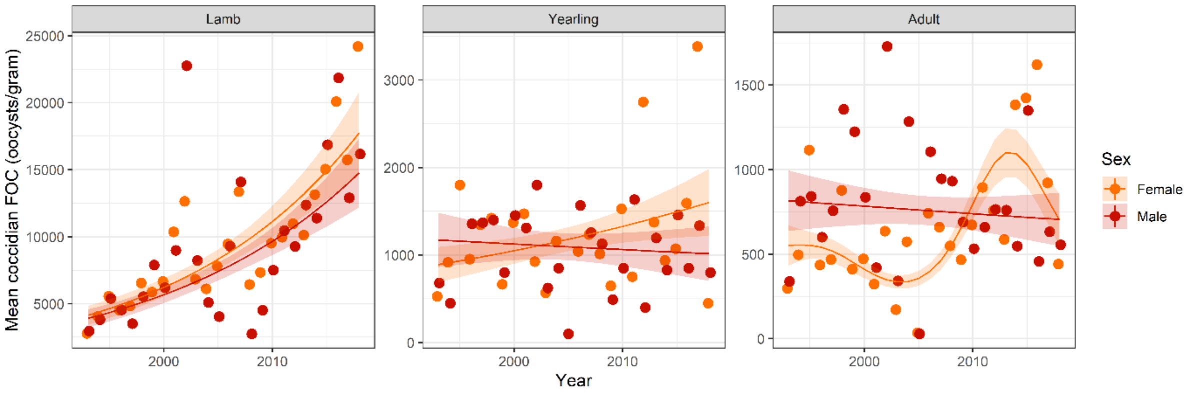


**Figure S4.** Changes in coccidian FOC across time as depicted in Figure 2B, but with y axes on different scales to emphasize the differences between sexes. Note the much higher FOC in lambs compared to the other two age classes, but also that the temporal patterns vary between the sexes in different ways in the three age categories, giving rise to the year-by-se-by-age interaction supported by model 11 in Table 3.

**Table S2.** A summary of linear models testing for variation in distributional parameters for strongyle FEC. 'Variable' indicates the parameter being analysed and 'Term' the variable being examined by the associated F-test and P-value.

| Model | Variable | Term | F | P |
|---|---|---|---|---|
| 1 | Variance | Year² | 0.17 | 0.684 |
| 2 | Variance | Year | 0.13 | 0.724 |
| 3 | Dispersion | Year² | 1.56 | 0.223 |
| 4 | Dispersion | Year | 0.04 | 0.851 |
| 5 | k | Year² | 3.81 | 0.061 |
| 6 | k | Year | 0.13 | 0.725 |
| 7 | Variance | 2 + Year:Age | 0.23 | 0.798 |
| 8 | Dispersion | 4 + Year:Age | 0.1 | 0.901 |
| 9 | k | 6 + Year:Age | 0.3 | 0.743 |
| 10 | Variance | 2 + Year:Sex | 0.01 | 0.925 |
| 11 | Dispersion | 4 + Year:Sex | 0.4 | 0.529 |
| 12 | k | 6 + Year:Sex | 0.06 | 0.811 |

**Table S3.** A summary of linear models testing for variation in distributional parameters for coccidian FOC. 'Variable' indicates the parameter being analysed and 'Term' the variable being examined by the associated F-test and P-value.

| Model | Variable | Term | F | P |
|---|---|---|---|---|
| 1 | Variance | Year² | 4.97 | 0.036 |
| 2 | Variance | Year | 27.69 | <0.001 |
| 3 | Dispersion | Year² | 0.08 | 0.778 |
| 4 | Dispersion | Year | 16.28 | <0.001 |
| 5 | k | Year² | 0 | 0.954 |
| 6 | k | Year | 0.05 | 0.824 |
| 7 | Variance | 1 + Year:Age | 12.39 | <0.001 |
| 8 | Dispersion | 4 + Year:Age | 0.56 | 0.575 |
| 9 | k | 6 + Year:Age | 5.21 | 0.008 |
| 10 | Variance | 1 + Year:Sex | 0.07 | 0.789 |
| 11 | Dispersion | 4 + Year:Sex | 1.92 | 0.172 |
| 12 | k | 6 + Year:Sex | 0.01 | 0.909 |

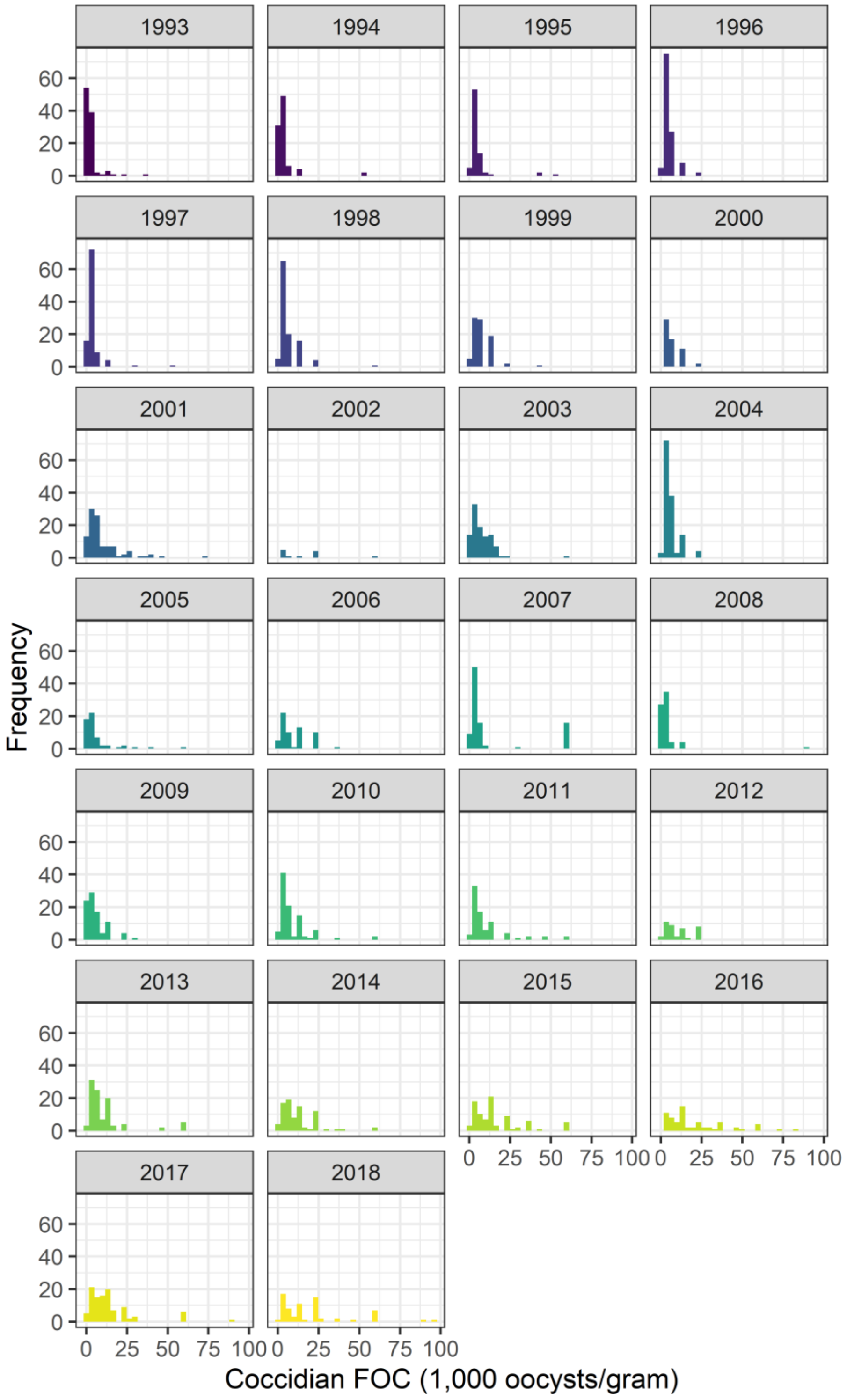


**Figure S5.** Histograms showing changes in the distribution of coccidian FOC across 26 years. In the early years, there is a more aggregated distribution with a greater proportion of animals having lower counts, while in later years more animals have higher counts and the distribution becomes les aggregated.
